# Ribosome biogenesis mediates the translational increase of non-optimal codon transcripts during IFN stimulation

**DOI:** 10.1101/2025.09.05.673799

**Authors:** Brenna N. Hay, Rachel Smid, Nathan Louie, Stephane Flibotte, Eric Jan, Leonard J. Foster

## Abstract

The interferon response is a signaling pathway unique to vertebrates that links the innate and adaptive immune responses. Interferons signal through a cascade of factors including the JAK-STAT pathway to induce the transcription of hundreds of interferon-stimulated genes (ISGs). Although the main interferon signal transduction pathways and ISGs have been elucidated, translational regulation of ISG transcripts is not fully understood. Prior work demonstrated that ribosomal protein RPL28 negatively regulates a subset of ISGs; however, we find that this effect may be due to a reduction in overall ribosome availability. Multi-omics analysis of RNA-seq and LC-MS/MS data reveal proteins, including several ISGs, that are translationally up-regulated in IFN-β-stimulated cells depleted of ribosome biogeneis factor BOP1. Analysis of codon usage demonstrates a significant reduction in codon optimality for proteins that are translationally up-regulated during BOP1 knockdown and IFN-β stimulation. Using reporter constructs, we demonstrate that codon non-optimal reporters are translated more than codon-optimized reporters in BOP1-depleted IFN-β cells. We propose that ribosome biogenesis regulates translational fine-tuning of integral protein production to ensure optimal interferon responses.

## Introduction

The interferon (IFN) response plays a key role in innate immunity, as it orchestrates signalling cascades that induce a rapid antiviral response to protect against pathogens (Le Bon and Tough, 2002; Wang *et al*., 2017). Upon detection of pathogen-associated molecular patterns (PAMPs) by pattern recognition receptors (PRRs), signaling phosphorylation cascades are activated resulting in the induction of key transcription factors, IRF3, to induce IFN production and secretion. IFNs, in a paracrine and autocrine manner, bind to their appropriate receptor to induce a second signalling cascade, resulting in the activation of transcription factors, STAT, to activate transcription of interferon-stimulated genes (ISGs). Proteins produced from the ISGs act on cellular machinery, leading to an anti-pathogenic state in the cell, with the ultimate goal of pathogen clearance.

As a crucial component of immunity, the IFN response must be finely tuned to prevent diseases that can arise from an over- or under-activating immune system (Melki and Frémond, 2020). For example, over-activation of IFN can lead to a range of autoinflammatory diseases, known as interferonopathies, which can induce varying effects within the central nervous system and the skin (Melki and Frémond, 2020). The IFN response is regulated by many negative feedback mechanisms. For instance, type I IFN receptor subunits are rapidly down-regulated upon the binding of a type I IFN to its receptor in order to prevent the same cell from re-initiating additional signaling (Constantinescu *et al*., 1995).

Recently, ribosomal proteins (RPs) have been implicated in translational controls of IFN signaling (Kerr *et al*., 2020). Reducing ribosome biogenesis has been shown to reduce mRNA levels of several immune-related transcripts including HMGB2, which produces a cGAS DNA-sensing protein, and IFNB1, the key cytokine in the IFN-β pathway and impacts virus infection by increasing HCMV replication (Bianco and Mohr, 2019). Conversely, ribosome biogenesis factors SBDS and SPATA5 were identified in a CRISPR loss-of-function screen, as they were required for flavivirus-encoded protein production (Ohlson *et al*., 2023). Ohlson and colleagues propose that a threshold ribosome abundance is required for viral protein expression for productive infection (Ohlson *et al*., 2023). Previously, we showed that during IFN-β stimulation in Hela cells, RPL28 was significantly increased in both the free ribosome and polysome fractions of purified ribosomes, suggesting a potential role for this RP in the IFN response (Kerr *et al*., 2020). RPL28 depletion resulted in pre-mature and enhanced production of ISGs in IFN-β-treated cells (Kerr *et al*., 2020). The link between ribosome biogenesis and translational regulation remains ambiguous, and the link to the host immune response remains to be investigated. In this study, we examine the relationship between ribosome biogenesis, translational controls and ISG expression. Our results point to codon usage as a parameter in regulating translation of mRNAs in the IFN response, thus providing an additional novel layer of control in IFN signaling.

## Materials and Methods

### Cell culture

A549 cells were cultured at 37°C in Dulbecco’s Modified Eagle’s Medium (DMEM, Gibco) supplemented with 10% fetal bovine serum and 1% penicillin-streptomycin.

### Trypan blue exclusion cell viability assay

Adherent A549 cells were aspirated, washed with phosphate buffered saline (PBS), and incubated at 37°C for 1 to 3 min with trypsin EDTA (Gibco) until cells began to lift, then mixed with DMEM to homogenize the cells. An equal volume of trypan blue (0.4%, Invitrogen) was added to 15 μL of the well-mixed cell suspension and gently mixed to prevent the formation of bubbles. A 10 μL sample was loaded onto a slide and inserted into an automated cell counter (Countess II, Thermo Fisher). Cell counts were performed using an optimized “A549” method which has been preset with cell size and shape settings specific to the cell line, while adjusting live and dead cell parameters with each slide as needed to ensure representative cell counts for all samples. Two-sample unpaired t-tests assuming equal variances were performed in Microsoft Excel on the cell viability obtained from three biological replicates.

### siRNA knockdown

A total of 3.0 x 10^5^ A549 cells were seeded into 6-well plates and incubated at 37°C for 24 h. Knockdowns were performed by adding 5 μL Dharmafect 1 (Dharmacon) to 195 μL Opti-MEM (Thermo Fisher) in one tube and 2.5 μL 20 μM target SMARTpool siRNA (Dharmacon) to 197.5 μL Opti-MEM in a second tube, then incubating at room temperature for 5 min. The contents of tubes A and B were combined and incubated at room temperature for 20 min. Media was aspirated from cells and replaced with this transfection mixture and 1.6 mL of antibiotic-free DMEM supplemented with 10% fetal bovine serum, allowing for incubation at 37°C for 48 h.

### Trifluoroethanol cell pellet lysis

Cells were washed with PBS, scraped for collection, and pelleted in a microcentrifuge at maximum speed for 15 min at 4°C. Any remaining PBS was removed by micropipette and the pellet was flash frozen in liquid nitrogen. Cells were resuspended by vortexing (VWR) with 150 μL of 50% trifluoroethanol (TFE) followed by sonication (Fisher Scientific) for 5 min in an ice bath. Samples were reduced and alkylated for 10 min at 95°C in a solution of 100mM Tris pH 8.5, 10 mM tris(2-carboxyethyl)phosphine (TCEP, Bioshop), and 40 mM chloroacetamide (CAA, Acros Organics). Samples were diluted to at most 10% TFE with 50 mM ammonium bicarbonate. Lys-C/trypsin (Promega) was added at a 1:50 ratio of protease:protein and incubated overnight at 37°C, followed by additional lys-C/trypsin at a 1:200 ratio incubated for 6 h at 37°C. Following TFE cell pellet lysis and digestion, peptide samples were desalted and purified on C-18 STAGE tips as described.(Rappsilber, Ishihama and Mann, 2003)

### LC-MS/MS (RPL28 knockdown)

Purified peptides were analyzed using a quadrupole time of flight mass spectrometer (Impact II; Bruker Daltonics) on-line, coupled to an Easy nano LC 1200 HPLC (Thermo Fisher) using nanoBooster with methanol and a Captive spray nanospray ionization source (Bruker Daltonics). Buffer A was composed of 0.1% formic acid and 2% acetonitrile in H_2_O and buffer B was composed of 0.1% formic acid and 80% acetonitrile in H_2_O. Samples were resuspended in buffer A and 100 ng were loaded onto the instrument running a standard 90-minute gradient. The Impact II was set to data-dependent acquisition.

### Protein identification and quantification (RPL28 knockdown)

A database search was performed using MaxQuant version 1.6.17 using label-free quantitation (LFQ).(Cox and Mann, 2008) Search parameters included the variable modifications methionine oxidation and N-terminal acetylation, and the fixed modification carbamidomethylation of cysteine residues. A 1% false discovery rate was applied at the protein and peptide level. Mass spectra were searched against a FASTA database for the *Homo sapiens (H. sapiens)* proteome from UniProt dated 2020.

### Proteomic statistical and bioinformatic analyses (RPL28 knockdown)

Bioinformatic analysis was performed in R using LFQ intensities for quantitation, and differential expression analysis was performed using *limma*.

### LC-MS/MS (BOP1 knockdown)

Purified peptides were analyzed using a quadrupole time of flight mass spectrometer (timsTOF Pro; Bruker Daltonics) operated in DIA-PASEF mode, coupled to an NanoElute UHPLC system (Bruker Daltonics) with Aurora Series Gen2 (CSI) analytical column (25 cm x 75 μM 1.6 μM FSC C18, with Gen2 nanoZero and CSI fitting; Ion Opticks, Parkville, Victoria, Australia). Buffer A was composed of 0.1% formic acid and 0.5% acetonitrile in H_2_O and buffer B was composed of 0.1% formic acid in acetonitrile. Samples were resuspended in buffer A and loaded onto the instrument running a standard 30-minute gradient.

### Protein identification and quantification (BOP1 knockdown)

A database search was performed using DIA-NN version 1.8.1 using label-free quantitation (LFQ) (Cox and Mann, 2008). Mass spectra were searched against a FASTA database for the *H. sapiens* proteome from UniProt. Search parameters included the fixed modifications N-terminal methionine excision and carbamidomethylation of cysteine residues, with no variable modifications. A 1% false discovery rate was applied at the protein and peptide level.

### Proteomic statistical and bioinformatic analyses (BOP1 knockdown)

Bioinformatic analysis was performed in RStudio using LFQ intensities for quantitation. Proteins were filtered to a minimum quantification of 25% across all samples. Filtered data was fitted to a linear model by limma (Ritchie *et al*., 2015). Gene ontology (GO) over-representation analyses were performed with the Panther web application (www.panterdb.org) using Fisher’s exact test and multiple testing correction of the p-value with the Benjamini-Hochberg procedure (FDR) (Mi *et al*., 2019; Thomas *et al*., 2022).

### RNA extraction

RNA samples were harvested with 500 μL TRI-zol (Invitrogen) and total cellular RNA was extracted according to the Direct-zol RNA Miniprep Kit, including the optional DNase I treatment (Zymo Research, R2052).

### RNA-seq

RNA samples were sequenced by the Sequencing Facility at the School of Biomedical Engineering (University of British Columbia) as follows. Sample quality control was performed using the Agilent 2100 Bioanalyzer or the Agilent 4200 Tapestation. Samples were prepared following the standard protocol for the Illumina Stranded mRNA prep (Illumina). Sequencing was performed on the Illumina NextSeq2000 with Paired End 59 bp x 59 bp reads. Sequencing data was de-multiplexed using Illumina’s BCL Convert. De-multiplexed read sequences were then aligned to the Homo sapiens (hg38 no Alts, with decoys) reference sequence using DRAGEN RNA app on Basespace Sequence Hub (Goyal *et al*., 2017).

### Transcriptomic statistical and bioinformatic analyses

Differential expression analysis was performed with DESeq2 in RStudio (Love, Huber and Anders, 2014).

### Codon usage bioinformatic analysis

APPRIS annotations and transcript sequences were downloaded from ensembl (www.ensembl.org) (release 112) (Harrison *et al*., 2024). Codon adaptation index (CAI) for each sequence was calculated using a locally installed version of the Perl script (CAIcal_ECAI_v1.4pl) (Puigbò, Bravo and Garcia-Vallve, 2008).

### Codon optimality constructs

The plasmids containing the extreme optimal and extreme non-optimal sequences were synthesized by Twist Biosciences into the pTwist CMV vector with insertion by HindIII and NheI. The extreme optimal and extreme non-optimal sequences were originally described and validated by Wu and colleagues; however, we included an ATG at the beginning of the extreme non-optimal region to initiate translation (Wu *et al*., 2019). The use of capital and lowercase letters separates the discrete components of the reporter, as outlined in each header.

T7-5’Bglo-ExOpt-GSGP2A-Nluc-3’Bglo AAGCTTTAATACGACTCACTATAAGGAGacatttgcttctgacacaactgtgttcactagcaacctcaaacagacaccatggccggcgataacggacccgaagatcgtgacaacggcgacgatggaggttatgctggaaagggagtcggaggcccaaaccctggaaacggcaccttccctggggggttctacggttattatggagccaagggggatttcgacatcgtcgctttcgggtactatggccgtcctatcggacctgggatcattcagaacttcgatgctgcttacgccgctgctatgccaattgagaaggaagatcccgctccatatattttccaggggggtaacgaaaagaacggaaccgctatcgtcggcgatgcaggaatggaaaaggatgactatggggaggaggtcgatcccgacccaatcatggatatgaacggtgagaccggggcatacaaggctgccgacgccggtacccgttatggtgaaatggaacccgctgccgaagatttcgccgacgaccaggagccaccagcctatgtcttcatcattaaggacatgcagggtccctattatgcagccaacttcggggaggacggtttcgaaggagctaaggatttcggcatgaccaacaccggcggtggtaaccgtgagatgaaggggtatgagttcgaacaggccgaagacggggaaaagcgtgaagaggaggagcctggcgacattaagtacatgggctatggtaacgccaaggctgccggaggccagattgagatggcaatgggcggtgcagggggatccggagctactaacttcagcctgctgaagcaggctggagacgtggaggagaaccctggacctAGCGGATCCAGGC CTATGGTGTTCACCTTAGAGGACTTCGTGGGGGACTGGAGGCAGACAGCAGGCTACAACCTGGACCAAGTGCTGGAGCAGGGAGGCGTAAGTTCACTATTCCAGAATCTGGGTGTCAGCGTCACACCCATCCAAAGGATTGTATTGAGCGGAGAAAATGGCCTGAAGATTGACATCCATGTCATCATTCCATATGAGGGGCTTTCCGGTGACCAGATGGGCCAGATTGAGAAGATCTTCAAGGTGGTCTACCCTGTGGATGACCACCACTTTAAAGTTATTCTCCATTATGGGACGCTGGTGATTGATGGAGTCACCCCCAACATGATCGACTACTTTGGCAGACCTTATGAAGGCATCGCTGTGTTTGATGGGAAGAAAATAACTGTTACTGGTACCCTGTGGAACGGGAATAAAATAATAGATGAGCGCCTCATCAACCCAGATGGTTCTTTGCTCTTCAGAGTGACTATCAATGGAGTTACAGGCTGGCGGCTTTGCGAGCGAATCCTGGCCTGAgctcgctttcttgctgtccaatttctattaaaggttcctttgttccctaagtccaactactaaactgggggatattatgaagggccttgagcatctggattctgcctaataaaaaacatttattttcattgcaa

T7-5’Bglo-ExNonOptwstartATG-GSGP2A-Nluc-3’Bglo AAGCTTTAATACGACTCACTATAAGGAGacatttgcttctgacacaactgtgttcactagcaacctcaaacagacaccatgcgaagtcaatcactgtcgacgtctctatcgctaaggcgcgtacgaagtttgaaacggagggcgcttctcccgctcagacgcaaaacagtatgcttttgtcatatattgaggcatccgacgagagtggttctttgcctcacaactgcgagatttcgatgtagcatatctgttgcgataaatagtaattgcgtatctttacatctgcaccataattcgtgggtttggttaaggcatagaagctcacacctgtgtcggccgagccatgtacaatctttacttcgctgtaaaacttctaggtcgaggctcctgcaaacgtccacgttatcatgctgtaattcacgatcgagactatgttcccatcttgtgtccaaatgcttaacactacaacgcaggctaataatactattgcaacttgtgactttgactagctcagtggttcggcacttgaattggacagttcaccaaactacgagtcatacatggctcactaataaatggacgttgcactctacaccgagctttactcgctcacgaccgacggcgttatccctttttacgagttccatacttgtatggtcttcccacctgtgcctcagactgctacgacaatttacacactcccgaagcgtgctacaccggctcaggatatgcgcgctgtcacggtggcgcgtaaaaatacgacaaagcagtccgttgagttgtagcacattagtaactagtggatccggagctactaacttcagcctgctgaagcaggctggagacgtggaggagaaccctggacctAGCGGATCCAGGCCTATGGTGTTCACCTTAGAGGACTTCGTGGGGGACTGGAGGCAGACAGCAGGCTACAACCTGGACCAAGTGCTGGAG CAGGGAGGCGTAAGTTCACTATTCCAGAATCTGGGTGTCAGCGTCACACCCATCCAAAGGATTGTATTGAGCGGAGAAAATGGCCTGAAGATTGACATCCATGTCATCATTCCATATGAGGGGCTTTCCGGTGACCAGATGGGCCAGATTGAGAAGATCTTCAAGGTGGTCTACCCTGTGGATGACCACCACTTTAAAGTTATTCTCCATTATGGGACGCTGGTGATTGATGGAGTCACCCCCAACATGATCGACTACTTTGGCAGACCTTATGAAGGCATCGCTGTGTTTGATGGGAAGAAAATAACTGTTACTGGTACCCTGTGGAACGGGAATAAAATAATAGATGAGCGCCTCATCAACCCAGATGGTTCTTTGCTCTTCAGAGTGACTATCAATGGAGTTACAGGCTGGCGGCTTTGCGAGCGAATCCTGGCCTGAgctcgctttcttgctgtccaatttctattaaaggttcctttgttccctaagtccaactactaaactgggggatattatgaagggccttgagcatctggattctgcctaataaaaaacatttattttcattgcaa T7-5’Bglo-Rluc-3’Bglo AAGCTTTAATACGACTCACTATAAGGAGacatttgcttctgacacaactgtgttcactagcaacctcaaacagacaccATGACTTCGAAAGTTTATGATCCAGAACAAAGGAAACGGATGATAACTGGTCCGCAGTGGTGGGCCAGATGTAAACAAATGAATGTTCTTGATTCATTTATTAATTATTATGATTCAGAAAAACATGCAGAAAATGCTGTTATTTTTTTACATGGTAACGCGGCCTCTTCTTATTTATGGCGACATGTTGTGCCACATATTGAGCCAGTAGCGCGGTGTATTATACCAGACCTTATTGGTATGGGCAAATCAGGCAAATCTGGTAATGGTTCTTATAGGTTACTTGATCATTACAAATATCTTACTGCATGGTTTGAACTTCTTAATTTACCAAAGAAGATCATTTTTGTCGGCCATGATTGGGGTGCTTGTTTGGCATTTCATTATAGCTATGAGCATCAAGATAAGATCAAAGCAATAGTTCACGCTGAAAGTGTAGTAGATGTGATTGAATCATGGGATGAATGGCCTGATATTGAAGAAGATATTGCGTTGATCAAATCTGAAGAAGGAGAAAAAATGGTTTTGGAGAATAACTTCTTCGTGGAAACCATGTTGCCATCAAAAATCATGAGAAAGTTAGAACCAGAAGAATTTGCAGCATATCTTGAACCATTCAAAGAGAAAGGTGAAGTTCGTCGTCCAACATTATCATGGCCTCGTGAAATCCCGTTAGTAAAAGGTGGTAAACCTGACGTT GTACAAATTGTTAGGAATTATAATGCTTATCTACGTGCAAGTGATGATTTACCAAAAATGTTTATTGAATCGGACCCAGGATTCTTTTCCAATGCTATTGTTGAAGGTGCCAAGAAGTTTCCTAATACTGAATTTGTCAAAGTAAAAGGTCTTCATTTTTCGCAAGAAGATGCACCTGATGAAATGGGAAAATATATCAAATCGTTCGTTGAGCGAGTTCTCAAAAATGAACAAGGGCgctcgctttcttgctgtccaatttctattaaaggttcctttgttccctaagtccaactactaaactgggggatattatgaagggccttgagcatctggattctgcctaataaaaaacatttattttcattgcaa

### RNA transfection

Cells were 60-80% confluent prior to RNA transfection. To create the transfection solution, 5 μL Lipofectamine 3000 (Thermo Fisher) was added to 195 μL Opti-MEM (Thermo Fisher) in one tube and 1 μg of capped and tailed RNA was added with Opti-MEM up to 200 μL in a second tube, then incubated at room temperature for 5 min. The contents of tubes A and B were combined and incubated at room temperature for 15 min. Media was aspirated from cells and replaced with this transfection mixture and 1.6 mL of antibiotic-free DMEM supplemented with 10% fetal bovine serum.

### Luciferase assay

Cells were washed with PBS and lysed in 500 μL 1X Passive Lysis Buffer for 15 min with gentle rocking (Promega). 25 μL of lysate was combined with 25 μL of the respective luciferase reagent (Promega; E2920, N1110) and immediately read on Spark microplate reader (Tecan).

### Sucrose gradient fractionation

Approximately 2×10^6^ A549 cells incubated in PBS with 0.1 mg/mL cycloheximide for 10 min at 37°C then lysed in mammalian lysis buffer (0.3 M NaCl, 15 mM Tris-Cl pH 7.5, 15 mM MgCl_2_, 1% Triton X-100, 0.1 mg/mL cycloheximide, 1 mg/mL heparin). Samples were centrifuged at 13000 rpm for 10 min at 4°C. The supernatant was then gently added to a 10-50% sucrose gradient. Gradients were spun at 35000 rpm at 4°C for 2 h 20 min in an ultracentrifuge (Beckman Coulter). Sucrose gradients were fractionated (Brandel) and the following settings were used for detection: sensitivity 0.5, chart speed 60, peak separator off, noise filter 1.5, manual, 0.375 mL/min. The surface area of the 60S peak was calculated in Fiji.(Schindelin *et al*., 2012) The total surface area was calculated by cutting out and weighing the trace.

## Results

### Ribosomal protein RPL28 depletion results in an increase in ISG proteins

We previously showed that depletion of RPL28 by siRNA transfection in IFN-β-treated Hela cells resulted in an increase in ISG expression (Wu *et al*., 2015; Kerr *et al*., 2020). We investigated whether this effect extends to other cells. We chose lung epithelial A549 cells as a model as the innate immune signaling pathway is relatively intact in these cells (Wu *et al*., 2015). Towards this, RPL28 was depleted by transfecting siRNAs in A549 cells for 48 h, followed by treating cells with IFN-β. Cells were harvested and lysed cells at 8, 12, and 24 h post-transfection and cell lysates were analyzed by LC-MS/MS. As controls, we also analyzed protein expression by LC-MS/MS of transfected cells with siRNAs targeting RPS26, RPS28, or an siRNA control targeting firefly luciferase. Both RPS26 and RPS28 have previously been implicated in the immune response (Cui *et al*., 2014; Gripp *et al*., 2014; Ferretti *et al*., 2017; Wei *et al*., 2019).

We first investigated whether depletion of RPL28 had effects on cell viability by using trypan blue cell staining at 24 and 48 h post-transfection. Cell viability was similar between RPL28 siRNA and control FLuc-siRNA transfected cells at 24 h post-transfection (83.3% compared to 82.3%, p-value = 0.29) and at 48 h post-transfection (74.3% compared to 73.3%, p-value = 0.37) (Fig. 1A). Thus, RPL28 knockdown did not have any obvious effects on cell viability in A549 cells.

**Figure 1.**
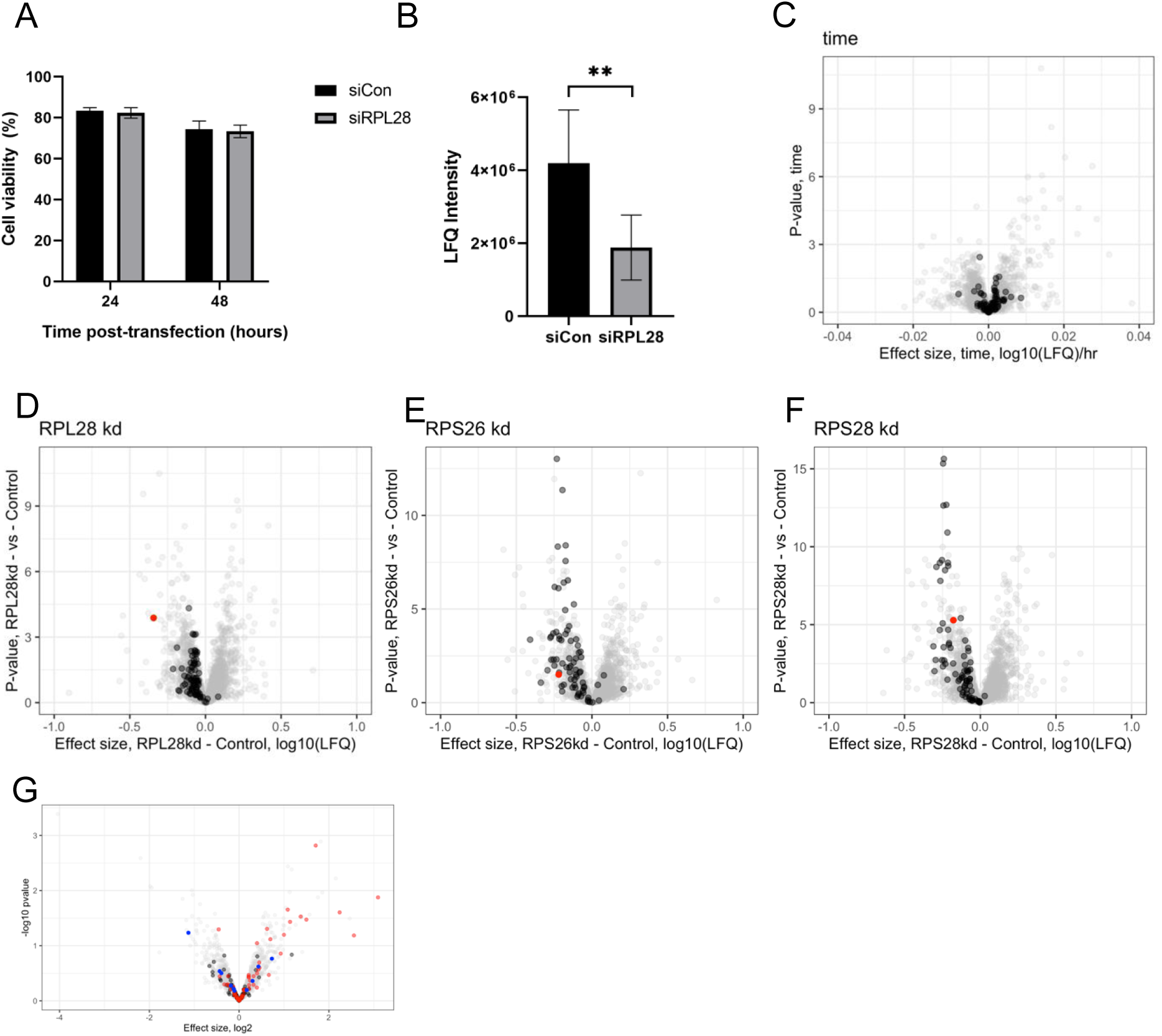
Targeted ribosomal protein knockdown results in general ribosomal protein depletion and ISG up-regulation during IFN stimulation. A) Percent cell viability of A549 cells transfected with RPL28 or control FLuc-siRNAs for the indicated times after transfection. Cell viability was measured by trypan blue staining and an unpaired t-test was performed. Error bars depict standard deviation. N=3. B) RPL28 protein levels as quantified by label-free quantitation (LFQ) intensities during siRNA knockdown on A549 cells as determined by mass spectrometry. T-test p-value = 0.0079. N=3. C) Differential expression of RPs (black) over time. N=3. D) Volcano plot showing differential expression of RPs (black) and the target of the siRNA knockdown, RPL28 (red), in RPL28 knockdown and control conditions at all time points. All other proteins are grey. N=3. E) Differential expression of RPs (black) and the target of the siRNA knockdown, RPS26 (red), in RPS26 knockdown and control conditions at all time points. All other proteins are grey. N=2. F) Differential expression of RPs (black) and the target of the siRNA knockdown, RPS28 (red), in RPS28 knockdown and control conditions at all time points. All other proteins are grey. N=3. G) Differential expression of RP depletion (resulting from RPL28, RPS26, and RPS28 knockdowns) on RPs (black), ribosome biogenesis proteins (blue), ISGs (red), and all other proteins (grey) on IFN-β-stimulated vs unstimulated A549 cells across all time points. N_expt_=8. N_control_=3.

We next used LFQ to examine whole-proteome changes in cells transfected with RPL28 siRNAs vs control-transfected cells. RPL28 was reduced by 37% in the knockdown condition as compared to control cells, thus validating the knockdown approach (p-value = 0.0079). To determine whether RPL28 depletion affects ribosome abundance as has been previously suggested (Robledo *et al*., 2008), differential expression analyses were performed across the three RP knockdown conditions (FLuc control vs RPL28, RPS26, RPS28) over time. Volcano plots depicted the targeted RPs across time as a control and across the RP knockdowns (Fig. 1C-F). Each individual RP knockdown showed a depletion of its target as expected (RPL28 37%; RPS26 29%; RPS28 18%), but other RPs were also reduced in each RP knockdown condition (Fig. 1G). Specifically, in RPL28 knockdown, 96% of the ribosomal proteins from both ribosomal subunits were downregulated (73/76 RPs) and 36% were significantly downregulated (27/76 RPs). In the RPS26 knockdown, 95.9% of ribosomal proteins were reduced (70/76 RPs), with 63% being significantly reduced (48/76 RPs). Furthermore, 99% of ribosomal proteins were reduced during RPS28 knockdown, with 68% being significantly reduced (52/76 RPs). We also found that several key ribosome biogenesis proteins were down-regulated. In contrast, the majority of ISGs were up-regulated upon RP knockdown, including several ISGs that were previously reported (Fig. 1G) (Kerr *et al*., 2020). These results demonstrated that depleting RPL28, RPS26 or RPS28 led to a broad reduction of ribosomal proteins from both subunits, thus suggesting that overall ribosome abundance is reduced.

### BOP1 knockdown results in an increase in ISG proteins

Next, we sought to identify whether ribosome biogenesis was the broader mechanism underlying the enhanced ISG expression in IFN-β-treated cells. Towards this, we depleted a key 60S subunit biogenesis factor, BOP1, through siRNA transfection. BOP1 is part of the PeBoW complex that is required for 60S maturation and processing of 28S and 5.8S rRNA (Strezoska, Pestov and Lau, 2000; Hölzel *et al*., 2005). A549 cells transfected with BOP1 or control FLuc siRNAs for 48 h were subsequently treated with or without IFN-β. RNA samples were collected at 8 and 12 h post-stimulation for analysis by RNA-seq while protein samples were collected at 8, 12, and 24 h post-stimulation for analysis by LC-MS/MS. Cells transfected with BOP1 siRNAs showed that BOP1 was depleted by 73% at the RNA level and 28% at the protein level (Fig. 2B,D). Furthermore, sucrose gradient fractionation of BOP1-depleted cells showed an average reduction of 19% in the 60S subunit (Supplemental Figure 1). Finally, cells transfected with BOP1 siRNAs did not adversely affect cell viability at 24 and 48 h under these conditions (Fig 2A). In summary, these results demonstrated BOP1 depletion in A549 cells that reduced 60S subunit levels.

**Figure 2.**
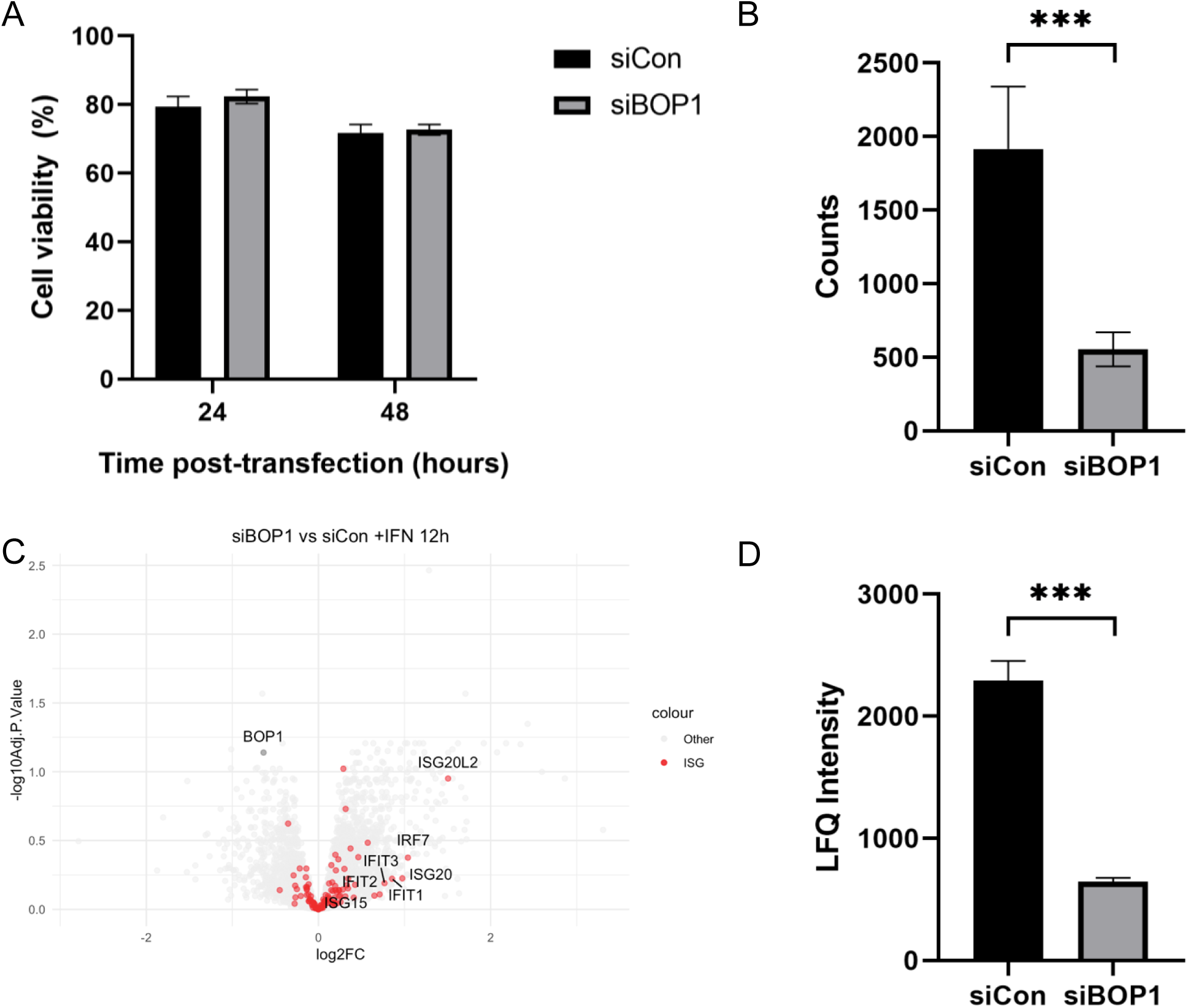
BOP1 knockdown results in ISG up-regulation during IFN stimulation. A) Percent cell viability of A549 cells transfected with BOP1 or control FLuc-siRNAs for the indicated times post-transfection. Cell viability was measured by trypan blue staining and an unpaired t-test was performed. Error bars depict standard deviation. N=3. B) BOP1 counts during siRNA knockdown on A549 cells as determined by RNA-seq. An unpaired t test was performed (p-value = 1.054E-7). N=3. C) Differential protein expression between A549 cells transfected with BOP1 or control FLuc-siRNAs with IFN-β stimulation for 12 h. ISGs are highlighted in red, BOP1 is highlighted in black, and all other proteins are in grey. N=3. D) BOP1 intensity during siRNA knockdown on A549 cells as determined by mass spectrometry. An unpaired t-test was performed (p-value = 1.685E-5). N=3.

We next analyzed the protein changes that occur between the control and BOP1 knockdown conditions in IFN-β-stimulated A549 cells. At 12 h post-IFN, a subset of ISGs (35/56 ISGs) was increased in BOP1-depleted cells compared to siFluc-transfected cells (Fig. 2C). We also identified several non-ISG proteins that were upregulated in BOP1-depleted cells including SARAF, GNB4, and ADCY9 (3296/6521). A one-sided Fisher’s exact test for enrichment of these subset of ISGs compared to non-ISGs amongst up-regulated proteins resulted in a p-value = 0.04918, demonstrating that these ISGs in particular are over-represented among the up-regulated proteins during BOP1 knockdown. These results showed that ribosome biogenesis results in an increase in proteins levels in a subset of ISGs under IFN treatment.

### A subset of proteins are translationally up-regulated during BOP1 knockdown

We next investigated whether the increase in ISG expression in IFN-β-treated BOP1-depleted cells is due to changes at the transcriptome level. Specifically, we focused on the subset of ISGs that were upregulated at the protein level (Fig 2C). RNA-seq analysis revealed minimal changes of the subset of ISG mRNAs, thus ruling out that transcription or mRNA decay effects on differential ISG protein expression (Fig. 3A).

**Figure 3.**
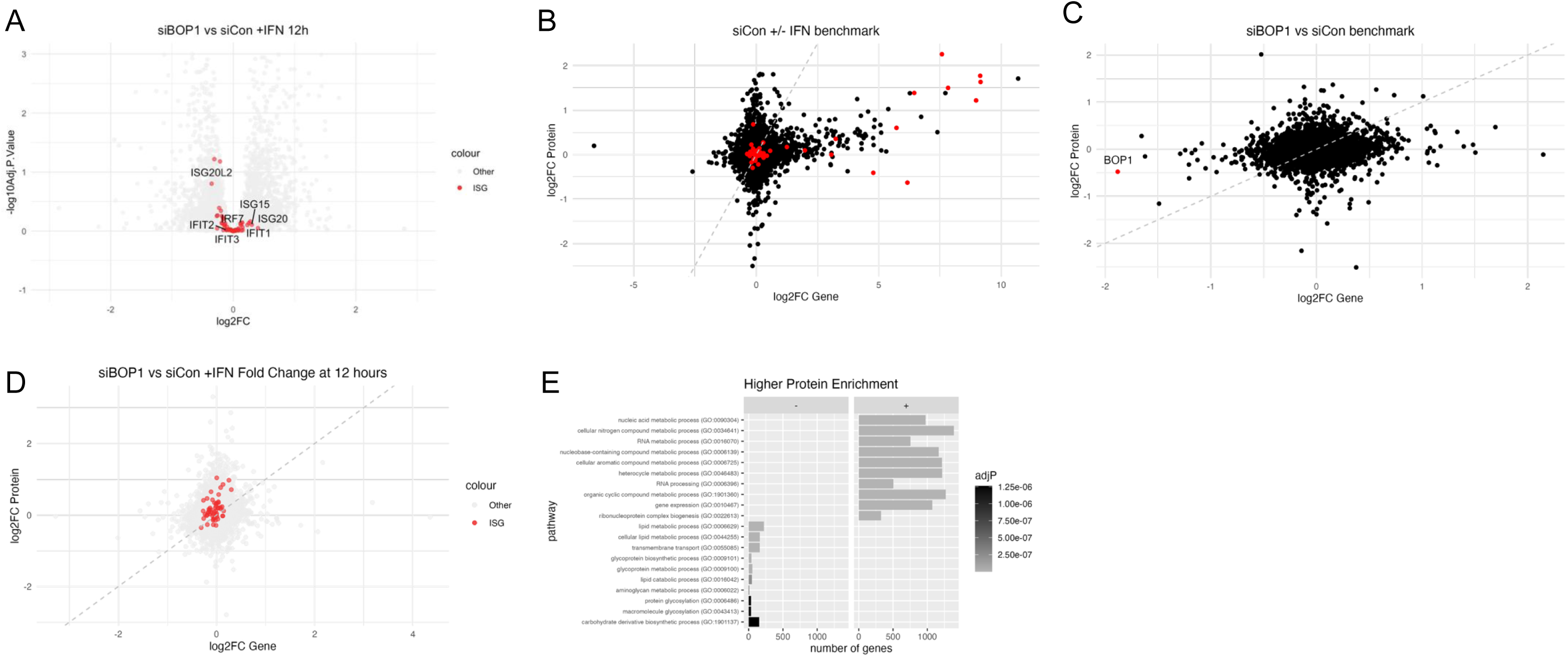
Multi-omics alignment reveals translationally up-regulated proteins. A) Transcript differential expression between A549 cells transfected with siBOP1 or the control siFLuc with IFN-β stimulation for 12 h. ISGs are highlighted in red and all other proteins are in grey. B) Alignment of differential expression in transcripts (x-axis) and proteins (y-axis) for A549 cells stimulated with IFN-β for 12 h during BOP1 knockdown compared to control FLuc knockdown. ISGs are highlighted in red, all other points are in grey. C) GO term analysis for terms up-regulated at the protein level. Signs (+ and -) depict up- or down-regulation respectively.

The transcriptome and proteome data suggested that the effects could be at the level of translation. We compiled both datasets from RNA-seq and LC-MS/MS and analyzed the fold-changes of protein and RNA to infer translational effects. First, we benchmarked this approach with the differential expression of A549 cells with and without 12 h of IFN-β stimulation (Fig. 3B) as well as in A549 cells transfected with BOP1 vs control siRNA (Fig. 3C). This first benchmark alignment revealed that several proteins and genes were up-regulated, many of which were ISGs, supporting that IFN stimulation results in an increase in the IFN pathway. A second benchmark was performed with the differential expression of A549 cells with BOP1 knockdown or a control. The second benchmark dataset demonstrated relatively normal distribution across all data points, while the profile of BOP1 also validated that while both the transcript and protein levels are reduced due to the siRNA knockdown (RNA 72.9% reduction; protein 28.3% reduction), there is a greater effect at the transcript level.

Next, we performed differential expression analysis of BOP1 vs control siRNA transfected cells at 12 h of IFN-β stimulation (Fig. 3D). Most ISGs showed an increase in differential expression at the protein level compared to the RNA level (41/54). Two ISGs, ISG15 and MX1, were previously identified by Kerr *et al* as part of a subset of ISGs that are upregulated during RPL28 knockdown (Kerr *et al*., 2020). These results strongly suggested that these ISGs are translationally upregulated in BOP1-depleted IFN-β-stimulated cells.

To determine whether other proteins were up-regulated at the protein level compared to the RNA level, GO term enrichment analysis was performed on the set of proteins that were up-regulated (Fig. 3E). GO terms involved in nucleic acid metabolism and RNA metabolism were enriched, while those involved in lipid metabolism were reduced. Subsequent analyses examine these datasets more closely.

### Proteins that are translationally up-regulated have lower codon optimality

Our results suggested that a subset of mRNAs is translationally upregulated in BOP1-depleted cells. One possibility is that there may be differences in codon usage of the ISG ORF that could impact translation. Codon usage of an mRNA is assessed by calculating the codon adaptation index (CAI), which quantifies how closely a gene’s codon usage matches the synonymous codon frequency within a reference set (Puigbò, Bravo and Garcia-Vallve, 2008). Using the tool CAIcal, the CAI was calculated for the aligned RNA-seq and LC-MS/MS datasets (Puigbò, Bravo and Garcia-Vallve, 2008). Using protein and mRNA datasets of IFN-beta cells (12 h) transfected with BOP1 or control siRNAs at 12 hours, the aligned datasets were then grouped by the CAI highlighted with distinct colours for all proteins (Fig. 4A) and for ISGs (Fig. 4C). The CAI values were also visualized in violin plots categorized by their differential expression at the protein or the RNA level (Fig. 4B,D). Amongst all proteins, those that were translationally up-regulated had a significantly lower codon optimality (0.778 vs 0.782, p-value = 0.0003576). A similar trend was observed amongst ISGs, though not statistically significant (0.772 vs 0.796, p-value = 0.1688). In summary, these results showed that proteins that exhibited greater differential expression at the protein level compared to the transcript level have reduced codon optimality as assessed by CAI.

**Figure 4.**
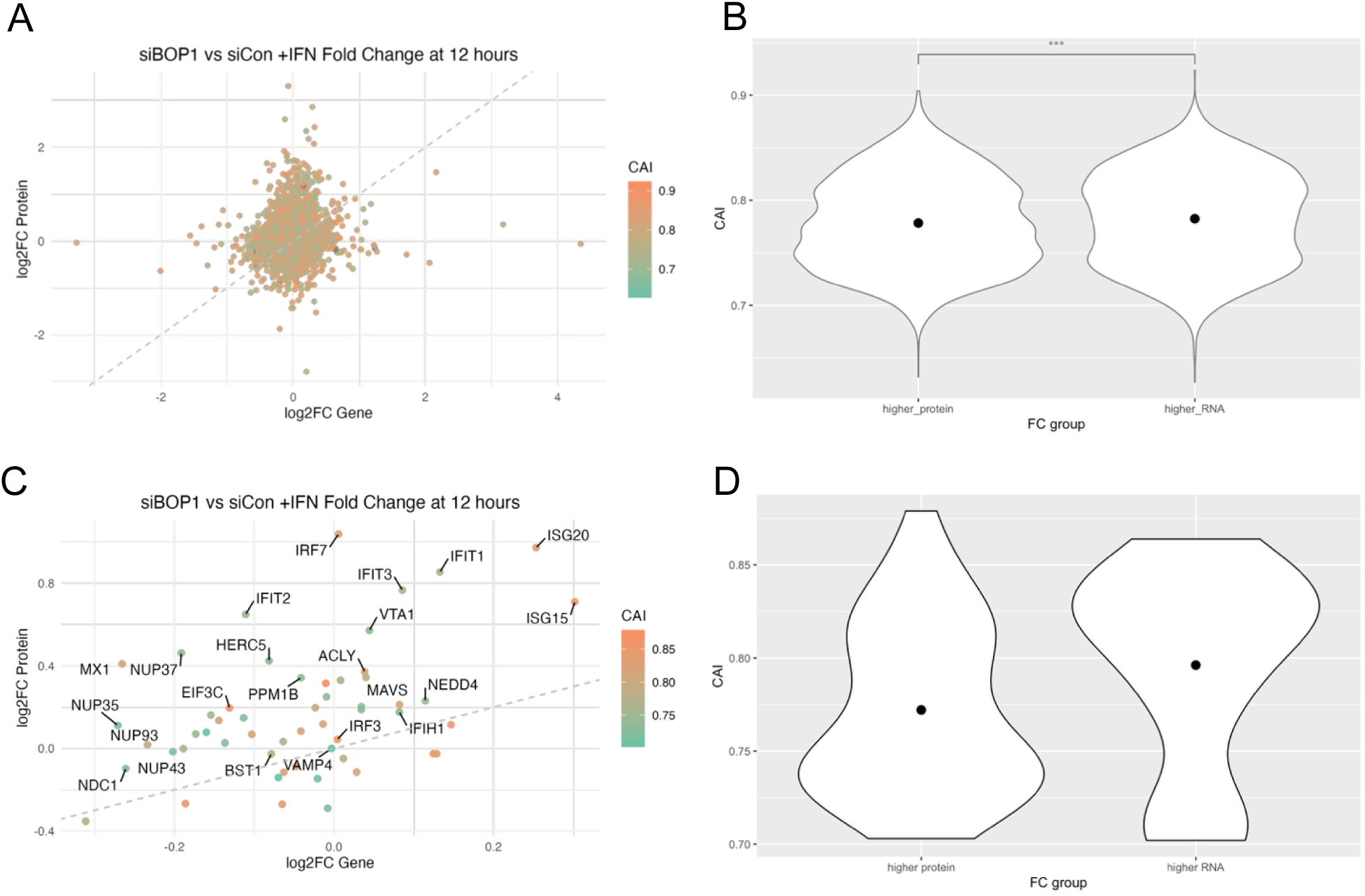
Translationally up-regulated proteins have lower codon optimality. A) Alignment of differential expression in transcripts (x-axis) and proteins (y-axis) for A549 cells stimulated with IFN-β for 12 h during BOP1 knockdown compared to control FLuc knockdown. All points are coloured by CAI. Grey dashed line depicts a slope of 1. B) CAI (y-axis) for differential expression results by fold change group (x-axis) including all proteins in dataset. Mean CAI of higher_protein = 0.778. Mean CAI of higher_RNA = 0.782. Unpaired t-test p = 0.0003576. C) Alignment of differential expression in transcripts (x-axis) and proteins (y-axis) for A549 cells stimulated with IFN-β for 12 h during BOP1 knockdown compared to control FLuc knockdown, plotting only ISGs. All points are coloured by CAI. Grey dashed line depicts a slope of 1. D) CAI (y-axis) for differential expression results by fold change group (x-axis) including all ISGs in dataset. Mean higher_protein = 0.772. Mean higher_RNA = 0.796. Unpaired t-test p = 0.1688.

### BOP1 knockdown results in an increase of translation of non-optimal transcripts

To determine whether codon optimality could dictate translation of an mRNA in BOP1 depleted cells, we monitored translation of extreme optimal (opt) and extreme non-optimal (non-opt) luciferase reporters adapted from Wu *et al* (Wu *et al*., 2019) (Fig. 5A). The optimal or non-optimal constructs consisted of a codon-optimized or -non-optimized region adapted from Wu *et al*, which was separated from a nanoluciferase reporter by a P2A sequence (Wu *et al*., 2019). The co-transfection construct contained only a Renilla luciferase reporter. All constructs were flanked by the 5’ and 3’ β-globin UTRs. In these experiments, BOP1 or control siRNA treated A549 cells stimulated with mock or IFN-β (2 h) were co-transfected with 5’capped and 3’poly-A tailed reporter mRNA (Fig. 5B). Cells were also co-transfected with an RLuc mRNA to control for transfection efficiency. Cells were harvested and luciferase activity was monitored. Relative NLuc activity was normalized to the RLuc co-transfection control to account for differences in global translation and transfection As expected, untreated cells transfected with the opt-Nluc reporter mRNA resulted in significantly more NLuc activity as compared to the non-opt-Luc reporter, demonstrating that the codon optimality can impact translation and stability of the mRNA (Fig. 5C). The mean fold change across each condition demonstrate a minimal increase in the translation of the optimal reporter under the control knockdown due to the IFN-β stimulation, and a decrease in the translation of the non-optimal reporter under the same conditions (Fig. 5D). Conversely, under the BOP1 knockdown condition, both the optimal and non-optimal reporters demonstrate increases in translation upon IFN-β stimulation. In order to assess the impact of the knockdown, we analyzed the mean fold change in the BOP1 knockdown compared to the control condition. In the unstimulated condition, there is little difference in the effect of the knockdown between the optimal and non-optimal constructs. In the IFN-β-stimulated condition however, the non-optimal construct exhibits a greater increase in translation due to the knockdown compared to the control (Fig. 5E). These results suggest that the BOP1 knockdown results in significantly greater translation of the non-optimal construct during IFN stimulation. Furthermore, to assess the impact of the IFN stimulation directly, we analyzed the mean fold change in the IFN-β-stimulated compared to the unstimulated condition. The effect of the IFN stimulation does not significantly impact the translation of the optimal construct between the control and BOP1 knockdown conditions. For the non-optimal construct, IFN stimulation reduced the translation in the control condition; however, IFN stimulation significantly increased the translation in the BOP1 knockdown condition (Fig. 5F). This work suggests that there is a specific and pronounced effect in which translation of non-optimal codons is increased during reductions in ribosome biogenesis, and is exacerbated during IFN-β stimulation.

**Figure 5.**
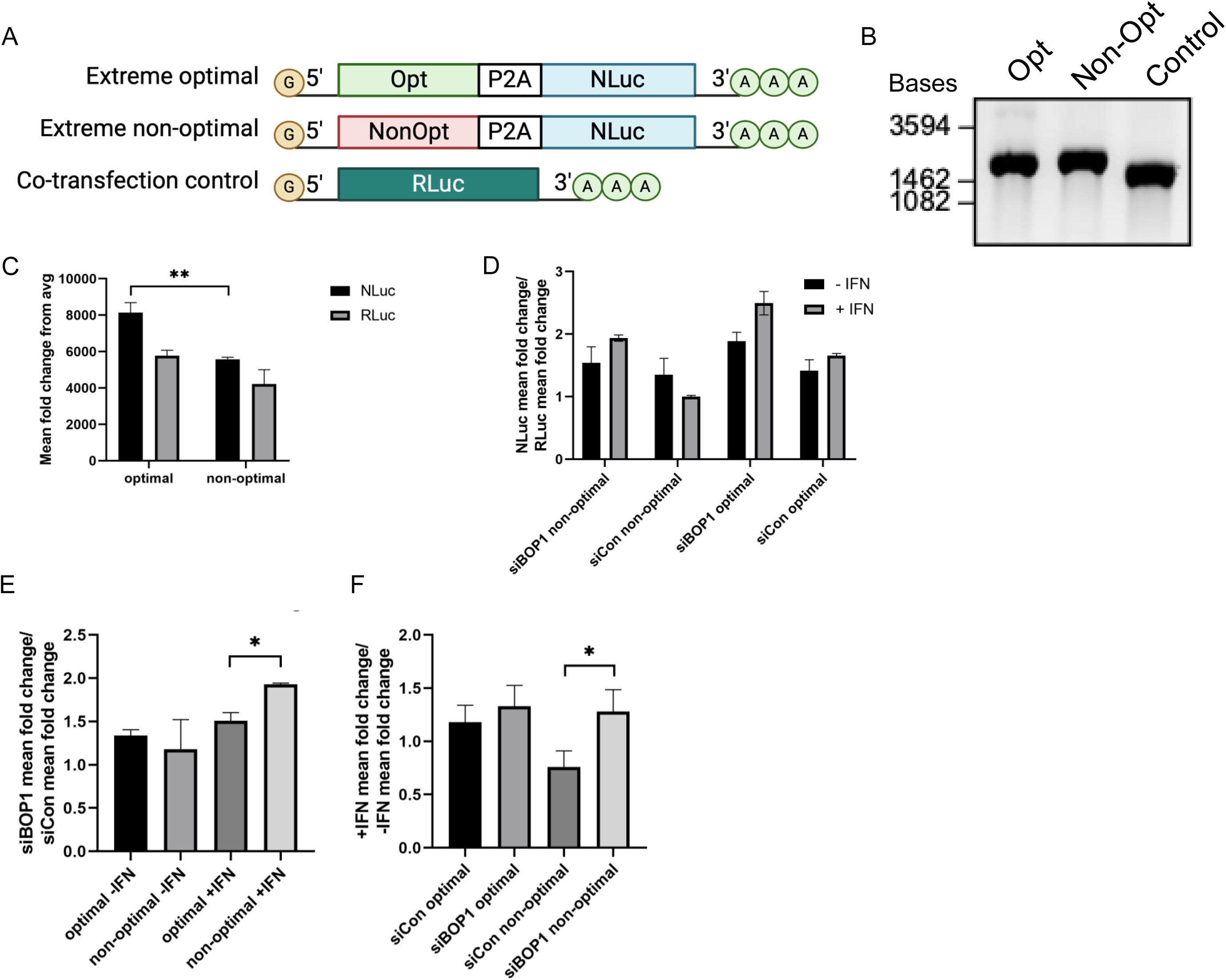
BOP1 knockdown increases translation of non-optimal transcripts under IFN stimulation. A) Codon optimality constructs adapted from Wu et al. (Wu *et al*., 2019). Extreme optimal (Opt) or extreme non-optimal (NonOpt) sequences were followed by P2A and nanoluciferase (NLuc), or a control with solely renilla luciferase (RLuc). All constructs were flanked by β-globin 5’ and 3’ UTRs. Created with BioRender. B) 1% RNA gel with 500 ng of each construct after capping and tailing. C) Mean fold change of NLuc and RLuc reporters in control knockdown condition with no IFN stimulation. P-value = 0.009. N=3. D) Mean fold change of NLuc over RLuc (y-axis) across each condition (x-axis). N=3. E) Relative luciferase activity normalized for the effect of BOP1 knockdown (y-axis) for each condition (x-axis). P-value = 0.01. N=3. F) Relative luciferase activity normalized for the effect of IFN-β stimulation (y-axis) for each condition (x-axis). P-value = 0.01. N=3.

## Discussion

The IFN response is a critical component of innate immunity and pathogen defense and as such requires dynamic gene and protein expression to ensure efficient defense without excessive inflammation and damage (Guillemin *et al*., 2022). Although this pathway is fundamentally initiated through changes in gene expression, multiple layers of post-transcriptional regulation exist to fine-tune the IFN response and include RNA binding proteins, mRNA stability, chemical modifications, and translational control (Burnett *et al*., 1998; Wu *et al*., 2019; McFadden *et al*., 2021; Rothamel *et al*., 2021; Guillemin *et al*., 2022). Many of these aspects of fine-tuning remain unclear and prompt further investigation.

Ribosome biogenesis is well-understood to play a key role in enabling effective translation; generally, when ribosome biogenesis is perturbed, this results in broad translational defects (Strezoska, Pestov and Lau, 2000; Kumar, 2021). Surprising roles for ribosome biogenesis have recently been identified, such as reductions in ribosome biogenesis increasing HCMV replication, which suggests the potential for nuance in the role of ribosome biogenesis in translational regulation, particularly in the immune response (Bianco and Mohr, 2019).

How does ribosome biogenesis impact the translation of the IFN response? The transcriptional landscape of the IFN response has been well-characterized across multiple species, including the identification of conserved ISGs (Mostafavi *et al*., 2016; Shaw *et al*., 2017). In contrast, there is less data supporting the changes in proteins that occur during IFN stimulation. Recent work investigating the proteome under IFN stimulation identified a translational lag of up to 24 h in the synthesis of ISG proteins, and suggest a potential role for ISGs that are produced earlier (Kerr *et al*., 2020). Further, a subset of ISGs was identified that were up-regulated upon RPL28 depletion, suggesting a role of the ribosome in negatively regulating the IFN response (Kerr *et al*., 2020). We sought to understand whether this effect may be more broadly connected to translation through ribosome biogenesis.

Depletion of BOP1, a key ribosome biogenesis factor, (Strezoska, Pestov and Lau, 2000) resulted in a significant up-regulation of ISGs after 12 h of IFN-β stimulation as compared to the knockdown control. This finding suggests that perhaps the observed effects on ISGs are a result of ribosome availability rather than composition. These effects are consistent to those observed during RP depletion with IFN-β stimulation (Kerr *et al*., 2020). This ISG up-regulation was demonstrated to be specifically occurring at the protein level, as no such effect was observed at the gene level. Although mRNA levels do not always result in equivalent protein levels, this finding pinpoints that the regulation is occurring post-transcriptionally (Schwanhäusser *et al*., 2011). The alignment of these data confirm that ISGs are translationally up-regulated during BOP1 knockdown and IFN stimulation, and furthermore provide a boon of information regarding levels of transcriptional and translational control that may be useful in understanding how ribosome biogenesis and the IFN response lead to changes in said control. We find that there is a small but significant difference in proteins that are translationally up-regulated having lower codon optimality.

GO term analysis on proteins that exhibited a greater increase in differential as compared to their transcripts identified several metabolic processes. The increase in these ranging metabolic processes suggest heightened cellular activity. Moreover, an enrichment in ribonucleoprotein complex biogenesis hints at increases in translation, which could relate to the translational shift observed during BOP1 knockdown. There was a decrease in GO terms associated with lipid metabolism, which could be a result of prioritization of metabolic processes during stress such as IFN stimulation. Although IFN stimulation does result in an increase in fatty acid oxidation and oxidative phosphorylation, these results were found after 24 h of IFN-α (Wu *et al*., 2016). Temporal changes during IFN stimulation are critical in understanding the dynamic nature of the IFN response and its impacts. This has been established as the changes in lipid metabolism shift over the course of IFN stimulation, where 6 h of IFN-γ stimulation resulted in reductions to triacylglycerol and lipid droplet levels, but 24 h of IFN-γ stimulation had the opposite effect (Truong *et al*., 2020). Inhibiting fatty acid synthesis has been shown to increase production of type I IFN, which also correlates with these findings (Kanno *et al*., 2021).

We next endeavored to identify how these ISGs may be translationally enhanced during ribosome biogenesis reduction. We identified a correlation between proteins that exhibit greater differential expression at the protein level compared to the transcript level and lower codon optimality. When testing codon optimality directly, our findings showed that non-opt reporter RNAs are increasingly translated during BOP1 knockdown and IFN-β stimulation. This work suggests that there is a specific and pronounced effect in which translation of non-optimal codons is increased during reductions in ribosome biogenesis, and significantly so during IFN-β stimulation. We propose that this effect is a form of translational regulation to fine-tune the immune response during times of stress, although this effect is likely more broadly applied across many critical pathways to ensure effective translation at appropriate times (Fig. 6). Under normal translation conditions, even optimal transcripts will experience some ribosomal stalling and possibly ribosome collisions and thereby may induce the ribosome quality control pathways, though the resulting proteins will be produced at a high level. Under BOP1 knockdown, there are fewer available ribosomes and therefore a concurrent reduction in stalling and collisions as well, resulting in a slight increase in translation of optimal transcripts. In contrast, under normal translation conditions, non-optimal transcripts can lead to greater stalling and ribosome collisions, which is known to result in mRNA instability and reduced protein production.(Wu *et al*., 2019) When BOP1 is knocked down, the reduction in ribosomes also reduces the stalling and collisions occurring on non-optimal transcripts, resulting in a translational up-regulation of non-optimal transcripts. In other words, when ribosome biogenesis is perturbed, this may result in a translational shift towards non-optimal transcripts, resulting in altered ratios of protein expression.

**Figure 6.**
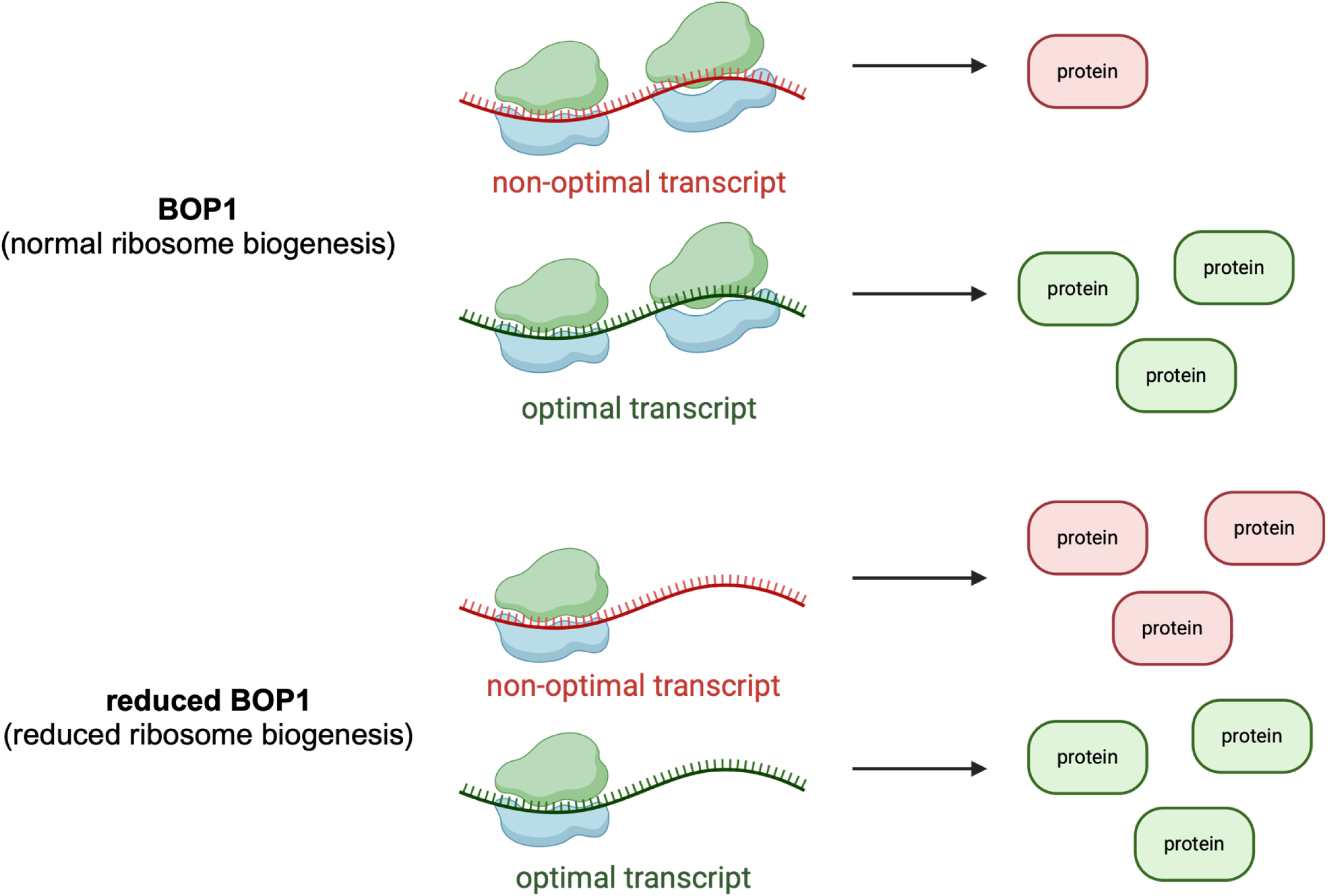
A proposed model illustrating the effects of BOP1 on codon-optimality-dependent translational regulation. Non-optimal transcripts and protein in red, optimal transcripts and protein in green. Created with BioRender. Under control conditions, BOP1 is present and ribosome biogenesis occurs. The translation of optimal transcripts is more efficient compared to non-optimal transcripts. Under conditions wherein BOP1 and therefore ribosome biogenesis is reduced, the ribosomal landscape shifts. Translation efficiency shifts and non-optimal transcripts are more efficiently translated.

Previously, Wu *et al*. demonstrated that the effect of codon optimality on transcript stability and protein production is dependent on layers translational regulation that may be occurring in cells, such as the translational shut-off that often occurs during virus infection.(Wu *et al*., 2019) During HSV-1 infection, they found that genes enriched in optimal codons were down-regulated and genes enriched in non-optimal codons were up-regulated. This work further supports our findings that during translational perturbations such as BOP1 knockdown, translation shifts towards non-optimal transcripts. Additional research is needed to further clarify the extent to which codon optimality impacts mRNA stability as opposed to translation efficiency.

These findings may impact the design of mRNA therapeutics, as one must consider the balance of CAI for both RNA stability and codon usage. mRNA therapeutics often employ codon optimization to enhance protein production by facilitating faster translation; however, if lower codon optimality can also promote effective translation, especially in the context of immune stresses such as the IFN response, over-optimization could be counterproductive, emphasizing the importance of tailoring mRNA therapeutics for specific conditions.

The experiment determining the effect of IFN-β stimulation on translational differences due to codon optimality was performed by performing a two-hour treatment prior to the mRNA transfection; however, to more closely replicate the earlier proteomics and transcriptomics experiments, IFN-β could be supplemented during the transfection as well. Further, to expand upon these results and pinpoint a specific mechanism, the effect of ribosome collisions in the observed results could be tested. BOP1 and ZNF598 could be knocked down, treated with IFN-β then co-transfected with mRNA for the extreme optimal or extreme non-optimal NLuc reporters and an RLuc reporter for analysis by luciferase assay. ZNF598 is a ubiquitin ligase which senses ribosome collisions and initiates the ribosome-associated quality control pathway, although a recent study identified that ribosome collision is not a strict requirement for ubiquitination by ZNF598 and that stalling alone can lead to degradation (Juszkiewicz *et al*., 2018; Miścicka *et al*., 2024). More broadly, to assess global differences in translation and ribosome loading during BOP1 knockdown, Ribo-seq could provide information on ribosome occupancy across a range of transcript codon optimality.

These findings provide a comprehensive dataset for the investigation of the effects of BOP1 knockdown during IFN-β stimulation, including establishing a transcriptome and translatome, as well as an alignment of differential expression providing insights on translational regulation. Notably, we observed that ISGs exhibited translational up-regulation during BOP1 knockdown and IFN-β stimulation. Upon analysis of the CAI for proteins that were translationally up-regulated and down-regulated, we found that all translationally up-regulated proteins have a small but significant reduction in CAI. ISGs that were translationally up-regulated shared this same trend with a greater difference in CAI, although not statistically significant. Functional analysis of the impacts of codon optimality revealed that BOP1 exerts an additional layer of translational control on codon optimality, which we propose is due to the availability of ribosomes impacting ribosome collision and degradation. We propose that this is a form of translational fine-tuning of the IFN response, as a means for the cell to produce protective proteins during times of cellular stress where translation machinery may be inhibited, but this requires further investigation on the functional mechanisms at play.

## Acknowledgements

The authors thank the Foster and Jan labs for their support, insight, and guidance, and particularly Jaden Chen for their assistance on the codon optimality work. This research was supported by the Natural Sciences and Engineering Research Council, Genome Canada, and Genome BC.

## Authorship

BNH, EJ, and LJF conceptualized this work and designed the experiments. BNH, RS, and NL performed experiments. SF performed the codon adaptation index analysis. BNH prepared the first manuscript draft; EJ and LJF edited and prepared the final draft with BNH.

## Data availability statement

The mass spectrometry data have been deposited to the ProteomeXchange Consortium via the PRIDE (Perez-Riverol *et al*., 2022) partner repository with the dataset identifier PXD060209.

The RNA-seq data have been deposited to the Gene Expression Omnibus (Edgar, 2002) with the dataset identifier GSE281848.

## Supplemental materials

**Supplementary figure 1.**

**The effect of BOP1 knockdown on the 60S fraction by sucrose gradient fractionation**. A) Sucrose gradient fractionation profiles during control (FLuc) and BOP1 knockdowns. N=2. B) Bar chart depicts the reduction in the 60S peak normalized to total surface area in the BOP1 knockdown condition compared to the control. Replicate 1 showed a 24% reduction in 60S during BOP1 knockdown and Replicate 2 showed a 14% reduction in 60S during BOP1 knockdown. N=2.

