## Supplementary figures and images for "Ribosome biogenesis mediates the translational increase of non-optimal codon transcripts during IFN stimulation"

### Supplemental Figure 1

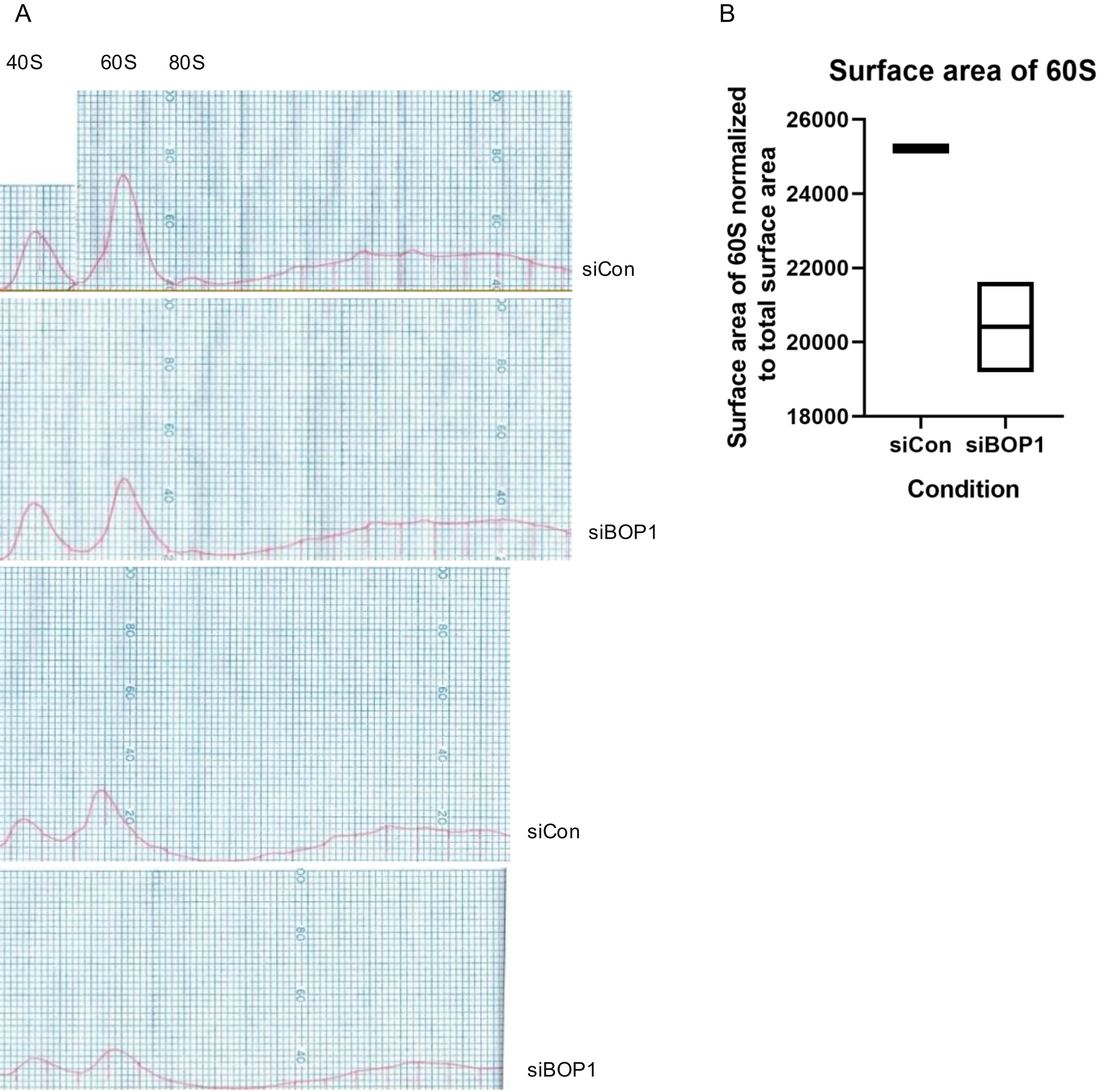
